# CNOT10 is involved in TTP-mediated AU-rich element containing mRNA metabolism, independent of mRNA decay regulation

**DOI:** 10.64898/2026.01.05.697706

**Authors:** Yukitomo Arao, Jason G. Williams, Wi Lai

## Abstract

Tristetraprolin (TTP)/ ZFP36 is an RNA binding protein that is involved in the turnover regulation of target adenylate-uridylate-rich RNA element (ARE) containing mRNAs. AREs are present in the 3’-untranslated region of transcripts expressed from many immediate-early genes, including cytokines and chemokines. It has been demonstrated that TTP-mediated post-transcriptional mRNA decay regulation is crucial for modulating physiological control, particularly in response to inflammatory stimulation. TTP is associated with the CCR4-NOT deadenylation complex through the TTP C-terminus to promote mRNA decay. However, it is not fully understood whether there are additional sites within TTP that contribute to its function through interacting with other factors. We analyzed the functionality of the unique tryptophan residues located in the TTP N-terminus using a cell-based assay system that consists of a tetracycline-responsive CMV promoter-driven, intron-inserted luciferase (LUC). This system enabled us to analyze TTP activity during both the early phase and the steady-state phase of gene expression, as well as in the post-transcriptional mRNA decay following the treatment of tetracycline analogs. Meanwhile, we identified putative TTP associates using a proximity labeling method. We found that tryptophan residues in the TTP N-terminus together with CNOT10, a component of the CCR4-NOT complex, were involved specifically in the reduction of the ARE-containing LUC mRNA level during the early phase of gene expression. However, they were not involved in the decay of LUC mRNA in the steady-state phase. We propose a novel post-transcriptional TTP functionality in the reduction of ARE-containing mRNA level, which differs from the well-characterized mRNA decay activity.

## Introduction

Tristetraprolin (TTP)/ ZFP36 is an RNA binding protein containing a C3H-type tandem zinc finger domain, which preferentially binds to the UUAUUUAUU RNA sequence known as the core motif of adenylate-uridylate-rich element (ARE) associated with mRNA turnover regulation (Carrick et al. 2004). AREs have been found in the 3’-untranslated region (UTR) of the mRNAs for various cytokines, chemokines, and several oncogenes. These genes are primarily categorized into immediate-early genes, and their transcripts are characterized as unstable mRNAs (Bakheet et al. 2001), resulting in rapid and transient expression in response to stimuli (Bahrami and Drablos 2016). In the conventional TTP knock-out (KO) mice, the extended half-lives of several ARE-containing mRNAs, such as *Tnf*, *Csf2* and *Ccl3*, were observed (Taylor et al. 1996; Carballo et al. 2000; Kang et al. 2011). The Increased TNF level due to the stabilization of its mRNA in the TTP KO mice resulted in a complex TTP KO phenotype, including weight gain failure and arthritis (Taylor et al. 1996). These findings suggested that TTP enhances the decay of ARE-containing mRNAs to maintain normal physiological responses. It has been shown that the C-terminal region of the TTP protein interacts with CNOT1, a fundamental component of the CCR4-NOT deadenylase complex, to facilitate the deadenylation of the poly(A) tail prior to mRNA decay (Fabian et al. 2013). CCR4-NOT complex consists of 10 components in the mammalian cells, including 3’-5’ exonuclease activity-containing units (CNOT6, 7 and 8) and an E3 ubiquitin ligase activity-containing unit (CNOT4). CNOT1 is a scaffold factor that assembles the components of the complex.

Thus far, other subunits (CNOT2, 3, 9, 10 and 11) are regarded as non-catalytic subunits (Shirai et al. 2014). We have demonstrated the physiological role of the TTP C-terminus, which is the only recognized domain that interacts with the CCR4-NOT complex, using the TTP C-terminal truncated knock-in (KI) mice (Lai et al. 2019). In comparison with the TTP KO mice, the TTP C-terminal truncated KI mice exhibited a less severe phenotype, characterized by occasional arthritis and a mild effect on *Tnf* mRNA stabilization in primary bone marrow derived macrophages (Lai et al. 2019). These results raised the possibility of additional interaction sites within the TTP protein or interacting factors that facilitate TTP function, beyond the interaction between the TTP C-terminus and CNOT1. However, it remains unclear whether there are additional interaction sites within the TTP protein or interacting factors that facilitate TTP functions. We investigated the potential role of the TTP N-terminus in post-transcriptional regulation, particularly the involvement of the unique tryptophan (W) residues (W32, W38, and W69). These tryptophan residues have been reported to interact with CNOT9, leading to the recruitment of TTP into the CCR4-NOT complex (Bulbrook et al. 2018). To assess the activity of TTP in post-transcriptional regulation, we developed a cell-based reporter assay system, which enabled us to analyze TTP functions in both the early phase and the steady-state phase of gene expression, as well as in the post-transcriptional mRNA decay. We also conducted a screening for TTP-associated factors using the proximity labeling method. We found that TTP is involved in the reduction of the ARE-containing mRNA accumulation during the early phase of gene expression. Additionally, CNOT10, a component of the CCR4-NOT complex, plays a role in this regulation by associating with the N-terminal tryptophan residues of TTP. However, the additive effect of CNOT10 was not observed in mRNA decay regulation during the steady-state phase. Our findings suggest that TTP has a novel functionality in the regulation of post-transcriptional gene expression.

## Results

*Assessing TTP activity on ARE-containing mRNA regulation using a Tet-ON/Tet-OFF reporter system* To analyze the TTP domain functionality, we developed a cell-based reporter assay system using the tetracycline-responsive CMV promoter-driven luciferase (LUC), which incorporated an intronic element in the LUC gene (TRE-INT-LUC) to define the expression levels of LUC pre-mRNA and mature LUC mRNA. A schematic diagram of the reporter gene and expressed products is illustrated in **Figure 1A**. The details of the reporter assay are described in Materials and Methods. Briefly, HEK293 cells were transfected with the reporter gene (TRE-INT-LUC) along with either Tet-ON or Tet-OFF expression plasmids to regulate the expression of TRE-INT-LUC and with or without TTP expression plasmid. The Tet-ON system enables the analysis of TTP activity at both the early gene expression phase, mimicking the acute expression of immediate-early genes, and the steady-state phase, as indicated by the LUC mRNA expression levels following treatment with a tetracycline analog, doxycycline (Dox). In contrast, the Tet-OFF system, which blocks TRE-INT-LUC expression through Dox treatment, can be utilized to analyze the effect of TTP on the post-transcriptional mRNA decay phase by monitoring the levels of LUC mRNA following Dox treatment.

**Figure 1.**
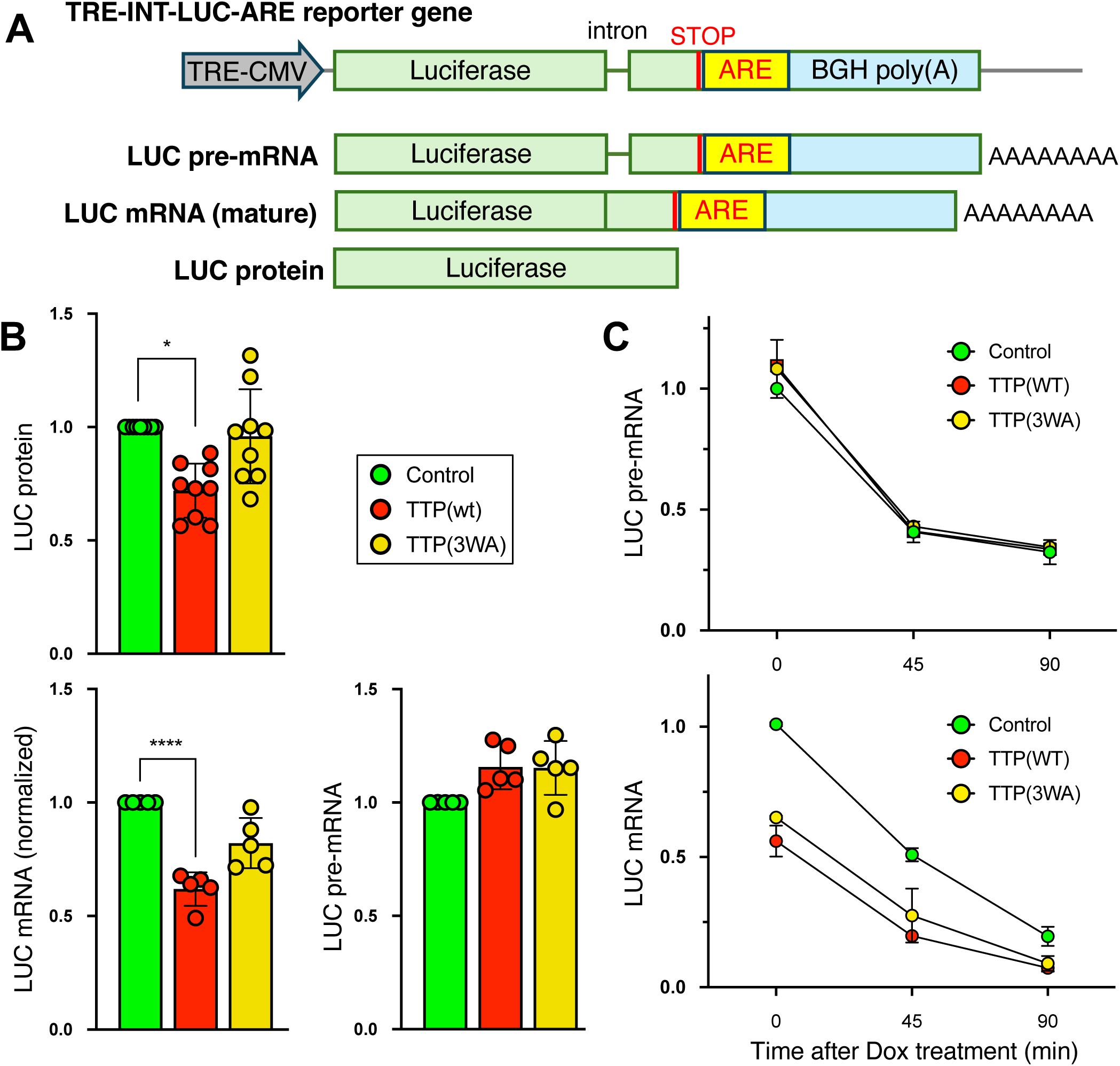
The effect of 3WA mutation of TTP on ARE-containing mRNA regulation. **A.** A diagram of the TRE-INT-LUC-ARE reporter gene is shown in the top panel. The products (LUC pre-mRNA, LUC mRNA, and LUC protein) from the reporter gene are illustrated. **B.** The samples were collected from the transfected cells after 6 hr of Dox treatment (Tet-ON assay). The relative levels of LUC protein are shown in the top panel (9 replicates). The relative levels of normalized LUC mRNA are shown in the bottom left panel (5 replicates), and the relative levels of LUC pre-mRNA are shown in the bottom right panel (5 replicates). The values from cells transfected with GFP and pcDNA3 (Control) were set as 1. The graph displays the values from replicated experiments and mean ± SD. Two-way ANOVA with Sidak’s multiple comparison test was performed to assess significant differences. * p < 0.05, **** p < 0.0001. **C.** The samples were collected from transfected cells after 45 min and 90 min of Dox treatment, as well as from non-treated wells for time 0 (Tet-OFF assay). The relative levels of LUC pre-mRNA are shown in the top panel. The relative levels of LUC mRNA are shown in the bottom panel. The results of Control (4 replicates), TTP WT (2 replicates) and TTP 3WA (2 replicates) are shown (mean ± SD). The values from cells transfected with GFP and pcDNA3 (Control) were set as 1.

We assessed the activity of GFP-fused full-length human TTP (TTP WT) and a mutated human TTP in which three tryptophan (W) residues in the N-terminus (W32, W38, and W69) were substituted with alanine (A) (TTP 3WA) on the ARE-containing mRNA regulation using a TRE-INT-LUC reporter containing 4 tandem UUAUUUAUU motifs (ARE) in the 3’ untranslated region (TRE-INT-LUC-ARE). First, we analyzed the effect of TTP WT and TTP 3WA on the LUC protein accumulation level expressed from the TRE-INT-LUC-ARE reporter gene after 6 hr of Dox treatment (Tet-ON). TTP WT lowered the LUC protein accumulation as expected. On the other hand, the 3WA mutation disrupted the TTP function (**Figure 1B**; top panel). Next, we assessed the LUC mRNA and LUC pre-mRNA levels at 6 hr after Dox treatment using quantitative PCR. The LUC mRNA level was presented as a normalized mRNA value, calculated as a ratio of the mature mRNA level to the pre-mRNA level. The profile of LUC mRNA levels (**Figure 1B**; bottom left panel) was similar to the profile of the LUC protein levels, as compared with the top panel of **Figure 1B**. Importantly, LUC pre-mRNA levels (**Figure 1B**; bottom right panel) were no different among these groups. These results suggested that the tryptophan residues in the N-terminus of TTP were involved in the lowering of the ARE-containing LUC mRNA level post-transcriptionally.

Furthermore, we analyzed the TTP function on the LUC mRNA decay after blocking the LUC gene expression with Dox treatment (Tet-OFF). Transfected cells were cultured overnight and then treated with Dox for 45 and 90 min before harvesting the cells for RNA extraction. We found that the LUC pre-mRNA level was reduced to the 45% level of non-treated cells within 45 min of Dox treatment, followed by a gradual reduction during the analysis (**Figure 1C**; top panel). The profiles of LUC pre-mRNA reduction were indistinguishable among the groups. In contrast, the steady-state LUC mRNA expression levels at time 0, a relatively long-term accumulation in LUC mRNA, were lower in both TTP WT and 3WA mutant expressing cells as compared to those in control (TTP non-expressing cells) (**Figure 1C**; bottom panel), suggesting that the 3WA mutation did not exhibit an altered effect on controlling the LUC mRNA level at the steady-state phase compared to that of WT. Moreover, we did not observe any significant differences in mRNA decay rate among the groups during chase of the LUC mRNA after Dox treatment, even between TTP WT and control (**Figure 1C**; bottom panel). These observations suggested that the TTP-mediated reduction of the reporter gene derived LUC mRNA was not due to an enhancement in mRNA decay.

### TTP associated CCR4-NOT complex components

We explored potential TTP-interacting factors using the TurboID-based proximity labeling (biotinylation) method to gain insights into the unexpected TTP activity observed in the reporter assay using TRE-INT-LUC-ARE. TurboID-fused WT TTP (TTP(wt)-TurboID) and C-terminal CNOT1 binding element truncated human TTP (TTP(ΔC)-TurboID) expression vectors were transfected into the HEK293 cells for screening. Biotinylated proteins were analyzed by mass spectrometry (MS). The abundances of putatively TTP-associating proteins (as determined by the sum of the areas under the chromatography peaks for each peptide mapped to that protein) were analyzed in TTP WT and ΔC transfected cells, with each group having five replicates. The fold-change levels were determined by comparing the abundance of peptides analyzed from nuclear export signal (NES)-fused TurboID transfected cells (five replicates). The experimental p-value was then calculated for each protein. When we applied the cutoff values of an experimental p-value < 0.01 and a greater than 2-fold change, 211 and 165 factors were identified from the TTP WT and ΔC expressing cells, respectively (**Supplemental Table S1.xlsx tab A** and **tab B**).

Although the truncation of the TTP C-terminus reduced the number of associated factors, 85.5% of TTP ΔC associates (141 factors) overlapped with those found in the TTP WT expressing cells, including CNOT1. This result implied that the C-terminus of TTP is unlikely to be the only surface interacting with the CCR4-NOT complex.

We examined the profiles of CCR4-NOT components in proximity labeling screening results (**Table 1**). Interestingly, there were no significant associations of the 3’-5’ exonuclease subunits of the CCR4-NOT complex, specifically CNOT7 and CNOT8 (CAF1 subunit), with either TTP WT or ΔC. On the other hand, there was a statistically significant association of CNOT6L (CCR4 subunit) with TTP WT but not with the ΔC mutant. The association of CNOT9 with TTP WT showed a lower fold-change (2.4x) and a lower consistency (p=0.017) compared to those with other subunits, such as CNOT2 (3.4x, p=4.8E-05), CNOT3 (3.7x, p=0.0028), CNOT10 (6.7x, p=3.2E-05) and CNOT11 (3.2x, p=0.0066). Noticeably, the C-terminal truncation of TTP resulted in the elimination of its association with CNOT9. CNOT7, CNOT8 and CNOT9 are known to be involved in shortening the poly(A) tail prior to mRNA decay (Bulbrook et al. 2018; Raisch et al. 2019). In contrast, we found that the association of CNOT10, whose function is not well understood, was significant with both TTP WT and ΔC, with the fold-change level comparable to that of CNOT1. We hypothesized that CNOT10 may play a role in the unexpected TTP activity described above, which differs from its function in promoting mRNA decay.

**Table 1.** Summary of fold-change levels and experimental p-values of CCR4-NOT components against TTP WT and TTP ΔC. The fold-change values over control (TurboID-NES) and the experimental p-values (Exp p-value) of CCR4-NOT components extracted from the cells transfected with either TTP(wt)-TurboID or TTP(ΔC)-TurboID (each group having five replicates) are shown. Italic values indicate statistically non-significant (the significant level was set as p < 0.01). nd indicates "p-value was indeterminable".

| | TTP WT | | TTP ΔC | |
| --- | --- | --- | --- | --- |
|  | Fold change | Exp p-value | Fold change | Exp p-value |
| CNOT1 | 7.9 | 1.0E-12 | 4.0 | 2.0E-10 |
| CNOT2 | 3.4 | 4.8E-05 | 3.3 | 2.6E-04 |
| CNOT3 | 3.7 | 2.8E-03 | 3.2 | 9.2E-03 |
| CNOT4 | *2.7* | *7.3E-02* | *2.8* | *1.1E-01* |
| CNOT6L | 4.1 | 6.6E-03 | *3.2* | *5.2E-02* |
| CNOT7 | *1.3* | *1.0E+00* | *1.1* | *9.9E-01* |
| CNOT8 | *2.1* | *4.8E-01* | *1.6* | *6.3E-01* |
| CNOT9 | *2.4* | *1.7E-02* | *0.4* | *nd* |
| CNOT10 | 6.7 | 3.2E-05 | 4.6 | 3.1E-04 |
| CNOT11 | 3.2 | 6.6E-03 | *1.5* | *1.3E-01* |

### Tryptophan residues in the TTP N-terminus are involved in the CNOT10 association

First, we performed a pull-down assay to assess the interaction between endogenous CNOT10 and the transfected GFP-fused TTP in HEK293 cells. However, we were unable to detect signals of endogenous CNOT10 in the pull-down assay using GFP-fused TTP (data not shown). Next, we employed the proximity labeling method to investigate the potential association between TTP and endogenous CNOT10. TurboID-fused WT and mutant TTPs were transfected into HEK293 cells. A schematic diagram of the fusion TTPs is illustrated in **Figure 2A**. The biotinylated endogenous CNOT10 was purified by streptavidin beads and then detected by Western blot with an anti-CNOT10 antibody. Similar levels of biotinylated CNOT10 were observed in cells that expressed TTP WT, 3WA, or ΔC. In contrast, the biotinylated endogenous CNOT10 was notably attenuated in the 3WAΔC expressing cells (**Figure 2B**), as compared to the expression levels of TurboID-fused TTPs shown in **Figure 2C**. To analyze the interaction between TTP and CNOT10 in more detail, we performed a pull-down assay using HEK293 cells that expressed GFP-TTP and FLAG-tag fused CNOT10 concurrently. We examined the interacting activity of CNOT10 with WT, ΔC, zfm, 3WA, and 3WAΔC TTPs. The cell lysates were purified with FLAG affinity beads. Then, the purified proteins were analyzed by Western blot with an anti-GFP antibody to detect GFP-fused TTP. Substantial levels of CNOT10 interaction with WT, ΔC, and zfm TTPs were observed. In contrast, the CNOT10 interaction was strongly attenuated by 3WA and 3WAΔC mutations (**Figure 2D**). The graph shows the normalized signal intensity of TTP-CNOT10 interaction against the input level (**Figure 2E**). These results suggested that the tryptophan residues in the N-terminus of TTP were crucial for CNOT10 interaction. Moreover, the observation of the TTP-mediated biotinylation of the endogenous CNOT10 (**Figure 2B**) likely indicated that both the N-terminus and C-terminus of TTP play a role in the interaction with CNOT10 within the CCR4-NOT complex.

**Figure 2.**
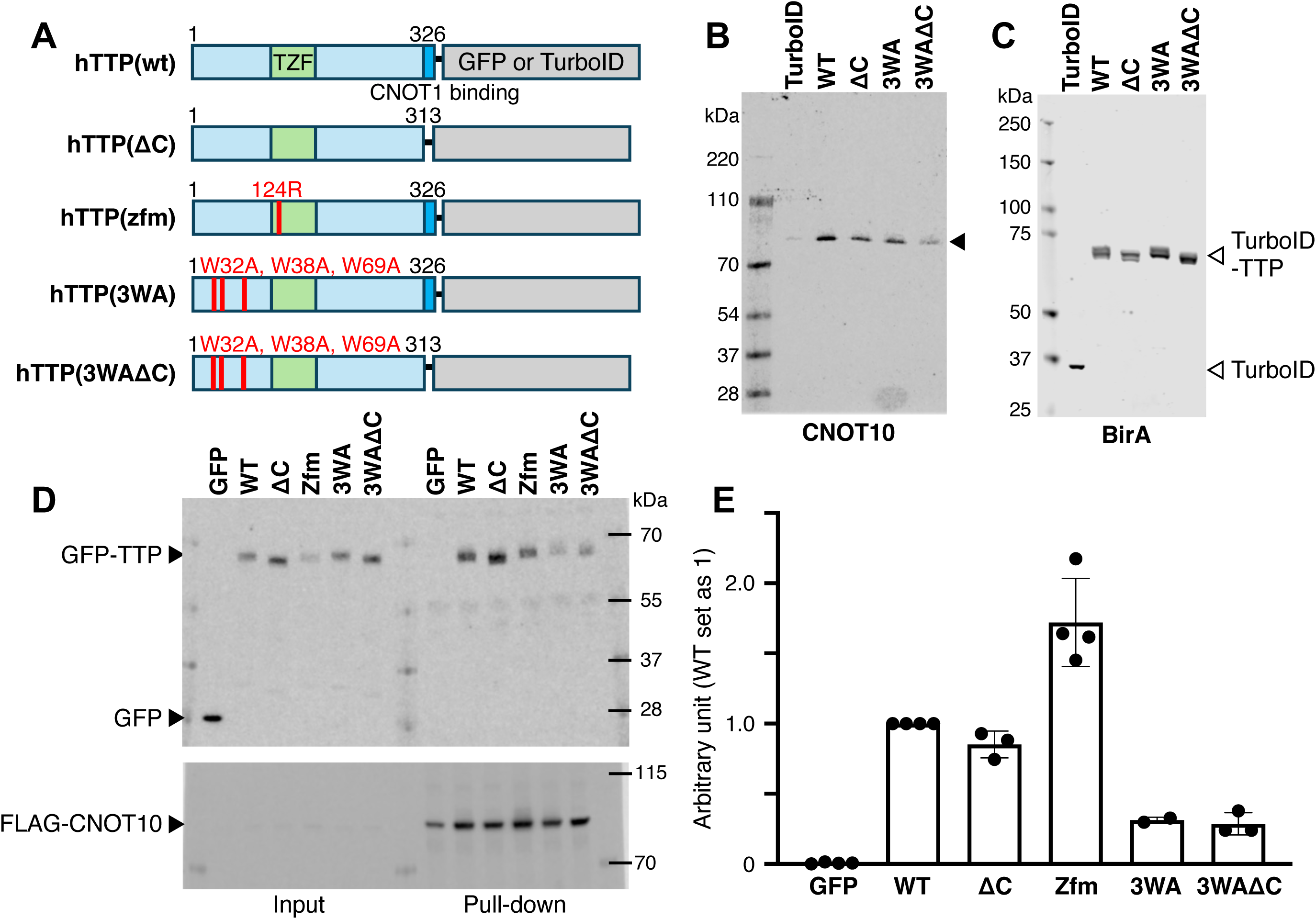
TTP-CNOT10 interaction. **A.** A diagram illustrates GFP- or TurboID-fused TTP WT and various TTP mutants. **B.** Cell lysates were prepared from the 4.5 hr of biotin-treated HEK293 cells that express TurboID-fused WT or mutant TTPs. Lysate from TurboID-NES (TurboID) expressing cells was used as a control. Biotinylated proteins were purified from the cell lysates by streptavidin magnetic beads. A representative Western blot shows the biotinylated endogenous CNOT10 using an anti-CNOT10 antibody. **C.** Representative Western blots show the levels of TurboID-NES and TurboID-fused TTPs in whole cell lysates using an anti-BirA antibody. **D.** Cell lysates were prepared from HEK293 cells that express FLAG-tag fused CNOT10 along with GFP-fused WT or mutant TTPs. FLAG-CNOT10 interacting proteins were collected with Anti-DYKDDDDK magnetic agarose. Representative Western blots show the pull-down GFP-fused TTP proteins using an anti-GFP antibody (top panel), and the levels of pull-down FLAG-tag fused CNOT10 using an anti-FLAG tag antibody (bottom panel). **E.** The relative level of signal density from pull-down samples, which was normalized to the input signal, is shown. The signal density from TTP(wt)-GFP was set as 1. The graph displays the values from 3-4 repeated experiments (mean ± SD).

### The effect of CNOT10 on TTP-mediated ARE-containing mRNA regulation

We used the TRE-INT-LUC-ARE reporter system to examine whether CNOT10 was involved in the TTP effect on the accumulation of ARE-containing mRNA. We included CNOT10 in a set of experiments similar to those described in the previous section (see **Figure 1B)**. The co-expression of CNOT10 enhanced the activity of TTP WT, resulting in a further reduction in the LUC protein levels compared to TTP WT with the control vector pcDNA3 (**Figure 3A**, top panel) after 6 hr of Dox treatment (Tet-ON).

**Figure 3.**
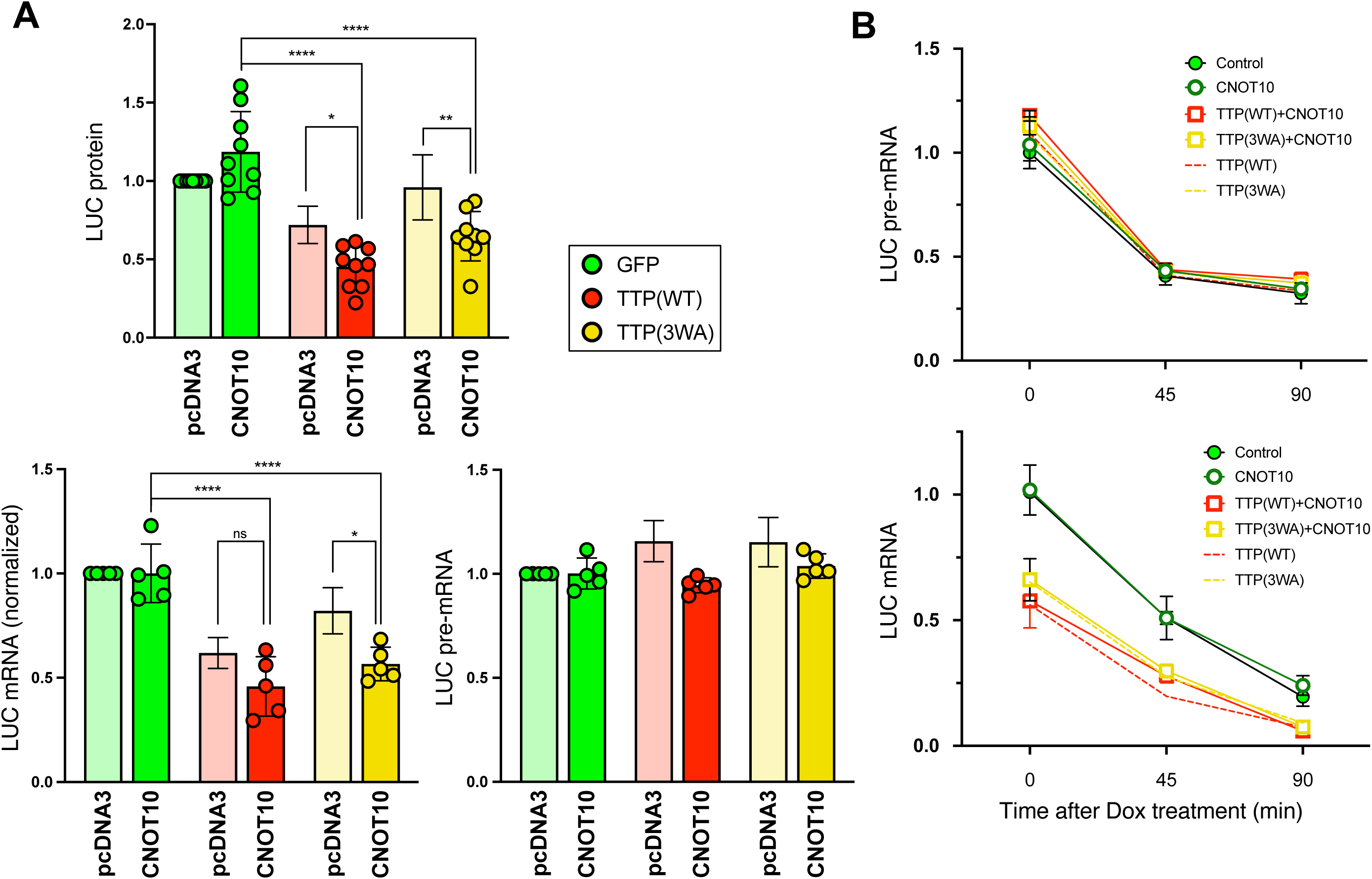
CNOT10 enhanced the TTP-mediated reduction of ARE-containing mRNA levels. **A.** Samples were collected from the transfected cells after Dox treatment for 6 hr (Tet-ON assay). The relative level of LUC protein is shown in the top panel (9 replicates). The relative level of normalized LUC mRNA is shown in the bottom left panel (5 replicates), and the relative level of LUC pre-mRNA is shown in the bottom right panel (5 replicates). The values from the cells transfected with GFP and pcDNA3 were set as 1. The graph shows the individual values from replicate experiments and mean ± SD for the CNOT10 co-expressed samples. TTP and pcDNA3 transfected samples are shown as pale-colored graphs with mean ± SD, which are reproduced from Figure 1. Two-way ANOVA with Sidak’s multiple comparison test was performed to evaluate significant differences. * p < 0.05, ** p < 0.01, **** p < 0.0001, *ns* no significance. **B.** The samples were collected from transfected cells after 45 min and 90 min of Dox treatment, as well as from non-treated wells for time 0 (Tet-OFF assay). The relative levels of LUC pre-mRNA are shown in the top panel. The relative levels of LUC mRNA are shown in the bottom panel. The results of Control (4 replicates), CNOT10 (4 replicates), TTP WT with CNOT10 (2 replicates) and TTP 3WA with CNOT10 (2 replicates) are shown (mean ± SD). The results for TTP WT and TTP 3WA are presented as dotted lines, which are replicated from Figure 1. The values from the cells transfected with GFP and pcDNA3 (Control) were set as 1.

Notably, the defect of the 3WA mutation in regulating LUC protein or mRNA levels was mitigated by the co-expression of CNOT10, which restored the TTP activity, leading to a reduction in both LUC protein and LUC mRNA levels (**Figure 3A**, top and bottom left panels). This suggested that the overexpressed CNOT10 may compensate for the weakened ability of the 3WA mutation in TTP to associate with the CCR4-NOT complex. The co-expression of CNOT10 with TTP did not affect LUC pre-mRNA levels (**Figure 3A**, bottom right panel), supporting that CNOT10-mediated enhancement of TTP function in lowering the LUC mRNA level was a post-transcriptional regulation. However, it should be noted that the co-expression of CNOT10 did not enhance the TTP-mediated LUC mRNA decay activity, as shown in **Figure 3B** bottom panel using the Tet-OFF system, when comparing the activity of TTP in the presence or absence of CNOT10 co-expression.

We analyzed the time course of LUC mRNA accumulation after Dox treatment using the Tet-ON system to evaluate the TTP function in the early phase of gene expression. RNA was extracted from the transfected cells at 2, 5, and 8 hr after Dox treatment. TTP WT decelerated the LUC mRNA accumulation compared to the control (**Figure 4**, top left panel). In contrast, TTP 3WA exhibited a LUC mRNA accumulation profile like that of the control (**Figure 4**, top right panel), indicating that the 3WA mutation diminished TTP function during the early phase of gene expression. Notably, the co-expression of CNOT10 significantly attenuated the accumulation of LUC mRNA with both TTP WT and 3WA (**Figure 4**, top panels). In contrast, the profile of LUC pre-mRNA accumulation in the co-expression of TTP and CNOT10 was found to be similar to that of the control (**Figure 4**, bottom panels).

**Figure 4.**
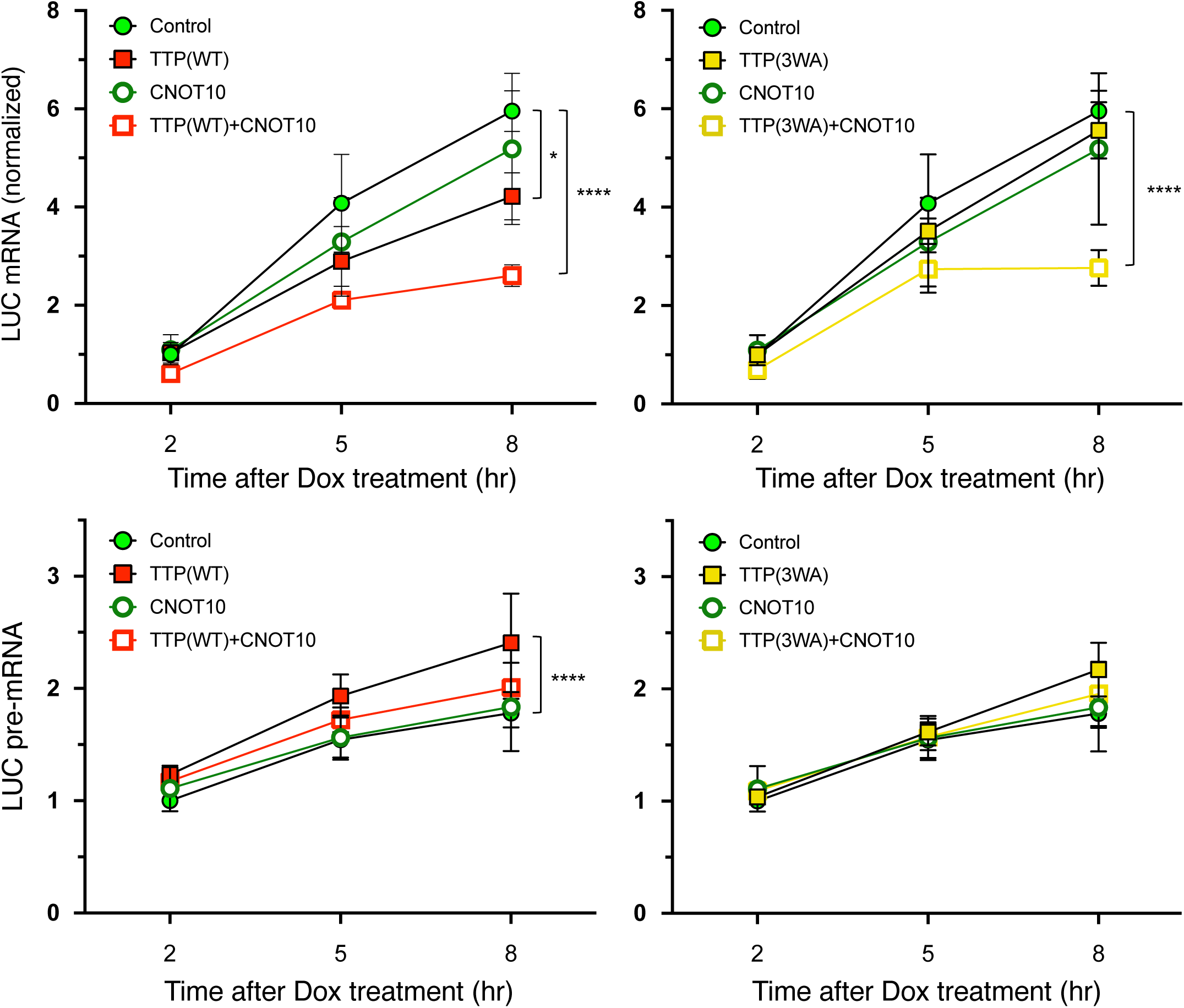
TTP-CNOT10 affects the early accumulation level of ARE-containing mRNA. RNA samples were collected from the transfected cells after 2, 5, and 8 hr Dox treatment (Tet-ON assay). The relative level of normalized LUC mRNA is shown in the top panels. The relative level of LUC pre-mRNA is shown in the bottom panels. The results (mean ± SD) from TTP WT (left; 3 replicates) or TTP 3WA (right; 3 replicates) transfected cells are shown. The graphs of GFP and pcDNA3 transfected cells (Control), and GFP and CNOT10 transfected cells (CNOT10) in the left and right panels are the same; the results from 6 experiments are shown. The values of the 2 hr time point for the Control were set to 1. Two-way ANOVA with Tukey’s multiple comparison test was performed to evaluate significant differences. * p < 0.05, **** p < 0.0001.

### TTP-CNOT10 complex

To understand the role of CNOT10 in TTP function, we conducted a pull-down assay combined with the proximity labeling method to identify factors associated with the TTP-CNOT10 complex. HEK293 cells were transfected with these two plasmids: TTP-TurboID (WT, ΔC or zfm) and FLAG tag-fused CNOT10. CNOT10 binding proteins were pulled down with FLAG tag affinity beads, and the harvested proteins were further purified with streptavidin beads. The collected proteins represent potential interactants of the TTP-CNOT10 complex, which were analyzed by MS. The abundance of peptides was analyzed in TTP WT, ΔC and zfm transfected cells, with each group in three replicates. The fold-change against TurboID-NES (three replicates) was assessed. The experimental p-value was then calculated. Potential factors of the TTP-CNOT10 complex (cutoff with experimental p-value < 0.01 and a greater than 2-fold change in WT) are shown in **Table 2**. The corresponding values for TTP ΔC and zfm are also included. Identified factors were few; however, CNOT1 and CNOT11 were listed as potential components of the TTP-CNOT10 complex. This result supported the validity of the screening. The association of CNOT1 and CNOT11 with the TTP-CNOT10 complex was reduced by C-terminal truncation of TTP, resulting in 65%, and 57% levels compared to TTP WT, respectively. Additionally, DDX3X, and RACK1 behaved similarly to CNOT1 and CNOT11, as their association was attenuated by C-terminal truncation of TTP in 65%, and 60% levels of TTP WT, respectively.

**Table 2.** Potential TTP-CNOT10 complex components. The fold-change values over control (TurboID-NES) and the experimental p-values (Exp p-value) of potential components of the TTP(WT)-CNOT10 complex are presented with a cutoff of experimental p-value < 0.01 and a greater than 2-fold change. The corresponding values for TTP ΔC and TTP zfm are also included (each group having three replicates). Additionally, the attenuation level associated with TTP ΔC is shown as % level compared to the TTP WT in the last column. Italic values indicate statistically non-significant. na indicates “it was not attenuated by TTP C-terminal truncation”

| ID | Symbol | TTP WT | | TTP ΔC | | TTP zfm | | Attenuation with ΔC (%) |
| --- | --- | --- | --- | --- | --- | --- | --- | --- |
|  |  | Fold Change | Exp p-value | Fold Change | Exp p-value | Fold Change | Exp p-value |  |
| O00571 | DDX3X | 32.9 | 5.05E-06 | 21.3 | 1.71E-05 | 30.6 | 4.19E-06 | 64.7 |
| A5YKK6 | CNOT1 | 13.4 | 1.25E-04 | 8.6 | 5.98E-04 | 14.9 | 8.41E-05 | 64.7 |
| O14654 | IRS4 | 6.5 | 2.33E-04 | 7.3 | 2.66E-04 | 10.1 | 9.68E-05 | na |
| D6REE5 | RACK1 | 74.1 | 4.92E-04 | 44.5 | 7.27E-04 | 55.0 | 9.07E-04 | 60.1 |
| H7C0C7 | CNOT11 | 29.7 | 6.97E-04 | 16.9 | 4.67E-03 | 51.0 | 7.40E-04 | 56.7 |
| Q9NZI8 | IGF2BP1 | 21.2 | 1.49E-03 | 26.7 | 6.79E-04 | *2.8* | *1.99E-01* | na |
| Q9Y5V3 | MAGED1 | 15.3 | 1.49E-03 | 17.7 | 1.48E-03 | 18.5 | 6.15E-04 | na |
| Q4VCS5 | AMOT | 23.2 | 2.84E-03 | 16.8 | 4.53E-03 | 32.9 | 2.92E-03 | 72.4 |
| Q9NQS1 | AVEN | 4.1 | 6.66E-03 | 2.7 | 2.35E-02 | 3.4 | 2.07E-02 | 66.7 |
| H0YKT5 | TLE3 | 3.9 | 8.01E-03 | 3.1 | 2.51E-02 | 3.2 | 3.31E-02 | 80.4 |
| D6RAA5 | TACC2 | 2.1 | 8.51E-03 | 1.6 | 3.70E-02 | *1.3* | *1.20E-01* | 79.3 |

## Discussion

CNOT10 is a relatively uncharacterized component of the CCR4-NOT complex. Initially, CNOT10 was identified as a new module of this complex, found alongside CNOT11/C2ORF29 in HEK293 cells stably expressing TAP-fused CNOT7 (Mauxion et al. 2013). Recent structural analyses suggested that the module of CNOT10 and CNOT11 complex is assembled into the CCR4-NOT complex through the N-terminal domain of CNOT1 (Mauxion et al. 2023; Levdansky et al. 2023). The function of CNOT10 in Trypanosoma brucei (Tb) has been reported, indicating that the depletion of TbCNOT10 caused the inhibition of mRNA turnover and deadenylation (Farber et al. 2013). Additionally, the report demonstrated that TbCNOT10 connected with the deadenylase TbCAF1 and the scaffold factor TbNOT1. However, in human cells (HEK293), CNOT10 was not necessarily required for the association of CAF1 (Caf1b; CNOT8) with CNOT1 (Farber et al. 2013). This finding aligns with another study that reported no change in the deadenylation rate in HEK293 cells with CNOT11/C2ORF29 knockdown (Mauxion et al. 2013). Furthermore, Lau et al. reported that CNOT6, CNOT6L and CNOT7 were not found in a pull-down with tagged CNOT10 from HeLa cells (Lau et al. 2009). In our experimental condition using HEK293 cells, the association of the CAF1 subunit (CNOT7 and CNOT8) with TTP was found to be insignificant. Because we used the proximity labeling method, we cannot exclude the possibility that the CAF1 subunit may be located at a distance from TTP within the CCR4-NOT complex, rather than being absent from the complex. On the other hand, the CCR4 subunit (CNOT6L) was associated with TTP WT in a statistically significant manner; however, it was not observed in the screening for the TTP-CNOT10 complex. Our observations were consistent with previous reports, which indicated that the 3’-5’ exonuclease subunits of the CCR4-NOT complex were not strongly associated with the CNOT10-CNOT11 module in mammalian cells. We also found that the TTP association with CNOT9 was less significant than that with CNOT10 under our experimental conditions. CNOT9 is known to be a non-catalytic component of the CCR4-NOT complex, and plays a role in mRNA decay regulation (Bulbrook et al. 2018; Pavanello et al. 2018). Additionally, tryptophan residues in the TTP protein have been reported to be involved in interactions with CNOT9, which is associated with TTP activity in ARE-mediated mRNA decay (Bulbrook et al. 2018). Our results suggest that TTP, which is associated with CNOT10 but lacks the CNOT9 binding in the CCR4-NOT complex, may play a distinct functional role in controlling ARE-mediated mRNA metabolism, which is different from the deadenylation of mRNA.

Recent reports suggest that the CCR4-NOT complex is involved in ribosome-associated quality control (RQC), which monitors stalled and collided ribosomes resulting from damaged mRNAs or transcripts with mutations, such as premature termination or non-stop mutations (D’Orazio and Green 2021; Muller et al. 2025). CCR4-NOT complex is recruited to the slowed or stalled ribosome via CNOT3, that is thought to be involved in the decay of mRNA (Zhu et al. 2024; Absmeier et al. 2023). We found RACK1 and DDX3X as potential components of the TTP-CNOT10 complex. RACK1 is known to be a ribosomal subunit and is directly involved in RQC; it cooperates with ZNF598, which ubiquitinates colliding ribosomes as a signal for RQC (Matsuo et al. 2017; Juszkiewicz et al. 2018).

RACK1 plays an essential role in positioning ZNF598 on stalled or colliding ribosomes (Sundaramoorthy et al. 2017). DDX3X plays multiple roles in post-transcriptional regulation, both in the nucleus and the cytoplasm, including RNA export and translation regulation (Soto-Rifo and Ohlmann 2013). However, it is unknown whether DDX3X is involved in RQC. Future research is needed to determine whether the TTP-CNOT10-mediated reduction of ARE-containing mRNA during the early phase of transcription is linked to the mRNA quality control coupled with translation. It has also been described that the nuclear function of the CCR4-NOT complex, which is involved in post-transcriptional RNA quality control and mRNA export from the nucleus in yeast (Azzouz et al. 2009; Assenholt et al. 2011; Kerr et al. 2011).

Notably, there are no analogs of CNOT10 and CNOT11 in yeast (Collart 2016). The nuclear functionality of the CCR4-NOT complex in mammalian cells remains poorly understood. Further investigations into nuclear post-transcriptional regulation via the CCR4-NOT complex in mammalian cells may be warranted to understand the TTP activity on the ARE-mediated mRNA metabolism.

We demonstrated that the unique tryptophan residues in the N-terminus of TTP are involved in reducing the accumulation of ARE-containing mRNA during the early phase of gene expression but not during the steady-state phase. Importantly, this effect differs from the well-known role of TTP in the ARE-mediated mRNA decay. TTP itself is recognized as an immediate-early gene that is activated by LPS or TNF in macrophages and monocyte cells (Carrick et al. 2004). It has been proposed that the increased TTP binds to ARE-containing mRNAs to promote their decay in the cytosol, thereby regulating the transient expression of immediate-early genes (Carrick et al. 2004). Our findings suggest a potential role for TTP in regulating the immediate-early gene expression by controlling the level of newly synthesized ARE-containing mRNAs. We anticipate that our observations will aid future research in a deeper understanding of TTP’s functionality in post-transcriptional gene regulation.

## Materials and Methods

### Plasmids

The plasmids hTTP(wt)-GFP, hTTP(ΔC)-GFP, and hTTP(zfm)-GFP were described in the previous report (Lai et al. 2019). The plasmids hTTP(3WA)-GFP and hTTP(3WAΔC)-GFP were made using the PCR primer-overlapping mutagenesis technique (Lai et al. 1998), using WT TTP or 1-313 (TTP ΔC) as a template. To generate the plasmids hTTP(wt)-TurboID, hTTP(ΔC)-TurboID, hTTP(3WA)-TurboID, and hTTP(3WAΔC)-TurboID, the TurboID coding element was amplified by Roche Expand High Fidelity PCR from the pcDNA3-V5-TurboID-NES (Branon et al. 2018) as a template with the following primers, TurboID_5’: 5-ACC GGT CGC CAA AGA CAA TAC TGT GCC TCT GAA GCT G-3’ and TurboID_3’: 5’-AGC GGC CGC TTT TCG GCA GAC CGC AGA CTG-3’. The amplified fragment was digested by NotI and AgeI. In the meantime, the plasmids of pBS-CMV-hTTP-EGFP (Lai et al. 2019) were digested with NotI and AgeI to remove the EGFP coding element. The TurboID coding element was swapped to the EGFP coding element of pBS-CMV-hTTP-EGFP plasmids, resulting in pBS-CMV-hTTP-TurboID. The plasmid pCMV6-hCNOT10 was obtained from Origene. To generate pGL3BGHn-TRE-INT-Luc, we obtained the plasmid CMV-LUC2CP/intron/ARE from Addgene (Younis et al. 2010).

The intron-containing LUC element was digested with NcoI and XbaI, then swapped into the NcoI and XbaI sites of the pGL3-TRE plasmid (Arao et al. 2004), resulting in pGL3BGHn-TRE-INT-Luc. To generate the TRE-INT-LUC-ARE plasmid, ARE-containing element of CMV-LUC2CP/intron/ARE was amplified by Roche Expand High Fidelity PCR with the following primers, Luc2CP-5: 5’-TTC TTC GAG GCT AAG GTG GTG GAC TTG G-3’ and Luc2CP-3: CGG CCG CCA CTA GTA AAT AAA TAA ATA AAT AAA TAA AAC T-3’. The element was digested with HpaI and SpeI, then swapped into the HpaI and XbaI sites of pGL3BGHn-TRE-INT-Luc. The correctness of all generated plasmids was confirmed by sequencing (GENEWIZ).

### Cell culture and transient transfection

HEK293 (human embryonic kidney, ATCC CRL-1573) cells were maintained in MEM + Earle’s salts with non-essential amino acids (NEAA) (10370-021; Gibco-Invitrogen) supplemented with 10% FBS (Hyclone), 4 mM L-glutamine (Gibco-Invitrogen) and 1% penicillin/streptomycin (Sigma). For transient transfections, the cells were cultured in MEM + Earle’s salts without NEAA (11090-081; Gibco-Invitrogen) supplemented with 10% charcoal-stripped FBS (Gemini-bio), 4 mM L-glutamine and 1% penicillin/streptomycin. The cells were seeded in 100-mm dishes at a density of 2 x 10^6^ cells per dish for the pull-down assay and the proximity labeling screening. The cells were transfected with various protein expression plasmids using Lipofectamine 2000 (Invitrogen) according to the manufacturer’s instructions. After 8 hr transfection, the medium (MEM + Earle’s salts without NEAA, containing 10% stripped FBS) was refreshed, and then the cells were cultured overnight.

### Tet reporter assay

HEK293 cells were cultured as described in the section on *Cell culture and transient transfection*. For the assay, cells were seeded in 24-well plates at a density of 0.9x10^5^ cells per well. The cells were transfected with the following DNA mixture for 8 hr using Lipofectamine 2000 (Invitrogen) according to the manufacturer’s instructions. The DNA mixture contained 10 ng expression plasmids for WT or mutant TTP, 100 ng expression plasmid for CNOT10, 400 ng doxycycline (Dox) responsive LUC expression plasmid (TRE-INT-LUC-ARE), and 200 ng Tet-On or Tet-Off expression plasmid (pTet-On, pTet-Off; Takara) was transfected into one well. After 8 hr transfection, the medium was refreshed, and then the cells were cultured overnight. The cells were treated with 200 ng/ml Dox for various time points before sample collection. The diagram of treatment schemes for each experiment is shown in **Supplemental Fig S1.pdf**. The cells were lysed with boiled 2x Laemmli buffer with DTT (100 µl/well) and then incubated at 90°C for 15 min to analyze the protein level. Twelve µl/sample was analyzed by Western blot using an anti-LUC antibody. Details are described in the section on *Western blot*. The cells were lysed with TRIzol (150 µl/well; Invitrogen) to analyze the RNA level. The cell lysates from the duplicate wells were combined to extract RNA (300 µl/sample). RNA extraction was performed following the manufacturer’s instructions.

### Quantitative PCR

The extracted total RNAs were treated with TURBO DNase (Invitrogen-Ambion) and purified with phenol-chloroform extraction. The purified RNAs were reverse transcribed using Superscript II (Invitrogen) and oligo dT16 (Invitrogen) to generate cDNA, following the manufacturer’s instructions. The quantitative PCR was performed using the QuantStudio 7 Flex (Applied Biosystems). The cDNAs were mixed with PowerUP CYBR Green Master Mix (Applied Biosystems) and the gene-specific forward and reverse primers for the reaction. The sequence of primers for Tet mRNA was pTet_F: 5’-CAT CAA GTC GCT AAA GAA GAA AGG GAA ACA-3’ and pTet_R: 5’-CGA TGG TAG ACC CGT AAT TGT TTT TCG T-3’; INT_F: 5’-GCT ACA AAC GCT CTC ATC GAC AAG GA-3’ and INT_R: 5’-GTC TCC TTA AAC CTG TCT TGT AAC CTT GAT-3’ for the LUC pre-mRNA; CDNA2_F: 5’-AAA TAC AAG GGC TAC CAG GTA GCC CCA GC-3’ and CDNA_R: 5’-CCA ACT TGC CGG TCA GTC CTT TAG GCA C-3’ for the mature LUC mRNA. The relative RNA level was calculated by a ΔΔCT model. To equalize transfection efficiency, the CT values for the mature mRNA and pre-mRNA of luciferase were normalized to the CT value of the Tet mRNA, which is expressed from the pTet-On or pTet-Off plasmid. To standardize expression variation in transiently transfected cell samples, normalized mRNA values were calculated as the ratio of mature LUC mRNA to LUC pre-mRNA.

### Western blot

Proteins were resolved by SDS-PAGE with 4-12% gradient gel and subsequently transferred to nitrocellulose membranes. The membranes were stained with Ponceau S before being subjected to Western blot analysis to verify the protein loading level. Blots were incubated overnight at 4°C with primary antibody for CNOT10 (1:600; A304-899A; Bethyl), GFP (1:1000; sc-8334; Santa Cruz), FLAG-tag (1:500; F1804; Millipore Sigma), luciferase (1:1000; EPR17789/ab185923; abcam), and BirA (1:1000; ab232732; abcam). The blots were washed and then incubated with IRDye 800CW-conjugated anti-rabbit antibody (1:20000; LI-COR Biosciences) for CNOT10, GFP and luciferase, or IRDye 800CW-conjugated anti-mouse antibody (1:20000; LI-COR Biosciences) for FLAG-tag and BirA. The signals were visualized by the Odyssey infrared imaging system (LI-COR Biosciences). The signals were quantified by using the Empiria Studio software (LI-COR Biosciences).

### Proximity labeling

The cells, which were transfected with TurboID-fused TTP expression plasmids (1 µg), were treated with 2 mg/l biotin for 4.5 hr, followed by an additional 1 hr incubation in fresh culture medium without biotin to reduce excess biotin. The cells were washed twice with PBS and then dissolved with boiled 2x Laemmli buffer (1 ml; BioRad) supplemented with 100 mM dithiothreitol (DTT). The samples were incubated at 90°C for 15 min and then stored at -20°C until use. To collect biotinylated proteins, the whole cell lysate suspended in Laemmli buffer was diluted to 4 ml in PBS, then gently agitated with streptavidin magnetic beads (NEB) at 25°C for 2 hr. The protein-bound beads were washed four times with PBS and then used for mass spectrometry analysis.

### Pull-down assay

After culturing overnight, the transfected cells with GFP-fused TTP expression plasmids (1 µg) and FLAG tag-fused CNOT10 expression plasmid (1 µg) were harvested, followed by a single wash with cold PBS, and then collected in 1ml of PBS into a 1.5 ml tube. The cells were precipitated by centrifugation at 4500 rcf at 4°C for 5 min. One ml Pierce IP Lysis buffer (Thermo Fisher Scientific) supplemented with protease/protein phosphatase inhibitor cocktail and EDTA (Pierce-Thermo Fisher Scientific) was added to the cells. The cell suspension was passed five times through a 25G syringe needle. The homogenate was centrifuged at 20800 rcf at 4°C for 15 min. The supernatant was collected into a new tube. 50 µl of supernatant was separated for evaluating the input level for the pull-down assay. Anti-DYKDDDDK magnetic agarose (Pierce-Thermo Fisher Scientific) was added to the remaining supernatant. The tube was placed on the shaking rocker at 4°C for 90 min. The protein-bound magnetic agarose was washed three times with wash buffer containing 50 mM Tris/HCl (pH 8), 150 mM NaCl, supplemented with protease/protein phosphatase inhibitor cocktail and EDTA (Pierce-Thermo Fisher Scientific). The protein-bound agarose beads were suspended in 2x Laemmli buffer with DTT, incubated at 90°C for 10 min and then stored at -20°C until use.

### Protein purification from TTP-TurboID and FLAG-CNOT10 co-expressed cells for MS analysis

After overnight culture, the cells transfected with the TurboID-fused TTP expression plasmid (0.5 µg) and the FLAG tag-fused CNOT10 expression plasmid (1 µg) were treated with 2 mg/l biotin for 4.5 hr, followed by an additional 1 hr in fresh culture medium without biotin. The cells were then harvested and lysed as described in the section on *pull-down assay*. After 20800 rcf centrifugation, the supernatant was collected into a new tube, and then the Anti-DYKDDDDK magnetic agarose was added. The protein-agarose complex was treated in the same way as described in the section on *pull-down assay*. The washed protein-agarose complex was suspended in 1 ml of 2x Laemmli buffer with DTT and incubated at 90°C for 10 min. The collected protein, suspended in Laemmli buffer, was diluted to 4 ml in PBS. The biotinylated proteins were purified and used for mass spectrometry analysis as described in the section on *Proximity labeling*.

### Mass spectrometry

Samples were prepared for mass spectrometric analyses using the EasyPep Mini MS Sample Prep Kit (Thermo Scientific) following the manufacturer’s protocol with modifications. 20 mg of modified trypsin (Promega) was suspended in 2 ml of 50 mM ammonium bicarbonate, pH 7.4 (100 µl), and added to each sample. Samples were digested overnight at 37 °C, shaking at 750 rpm. After the centrifugation for 3 min at 1500 rcf, the supernatant was collected into a new tube. The pellet (bead) was washed with 50 µl of 50 mM ammonium bicarbonate, pH 7.4; the supernatant was combined with the digested sample.

Digests were reduced and alkylated by the addition of 50 µl of reduction solution and 50 µl of the alkylation solution from the EasyPep Mini MS Sample Prep Kit. Samples were incubated for 10 minutes at 95 °C, and then 50 µl of the digest stop solution was added after the samples had cooled. Peptide clean-up was performed using the manufacturer’s recommended protocol for the EasyPep Mini MS Sample Pep Kit. Protein digests were analyzed by LC-MS/MS on a Q Exactive Plus mass spectrometer (Thermo Fisher Scientific) interfaced with a nanoAcquity UPLC system (Waters Corporation) equipped with a 75 mm x 200 mm HSS T3 C18 column (1.8 mm particle, Waters Corporation) and a Symmetry C18 trapping column (180 mm × 20 mm) with 5 mm particle size (Waters Corporation) at a flow rate of 450 nl/min. The trapping column was positioned in line of the analytical column and upstream of a micro-tee union, which was used both as a vent for trapping and as a liquid junction. Trapping was performed using the initial solvent composition. Approximately 1 mg of peptide digest was injected onto the column. Peptides were eluted by using a linear gradient from 99% solvent A (0.1% formic acid in water (v/v)) and 1% solvent B (0.1% formic acid in acetonitrile (v/v)) to 40% solvent B over 100 min.

For the mass spectrometry, a data-dependent acquisition method was employed with a dynamic exclusion time of 15 sec and the exclusion of singly charged ions. The mass spectrometer was equipped with a NanoFlex source and a stainless-steel needle and was used in the positive ion mode. Instrument parameters were as follows: sheath gas, 0; auxiliary gas, 0; sweep gas, 0; spray voltage, 2.7 kV; capillary temperature, 275 °C; S-lens, 60; scan range (m/z) of 375 to 1500; 1.6 m/z isolation window; resolution: 70,000; automated gain control (AGC), 3 × 10e6 ions; and a maximum IT of 100 ms. For the MS/MS scans: TopN: 10; resolution: 17500; AGC 5 x10e4; maximum IT of 50 ms; and an (N)CE: 27. Mass calibration was performed before data acquisition using the Pierce LTQ Velos Positive Ion Calibration mixture (Thermo Fisher Scientific). Data were processed in Proteome Discoverer (Thermo Fisher) using a workflow that included nodes for Minora Feature Detector, Sequest HT, and Percolator. Data were searched against the appropriate database using Sequest and included settings of: trypsin specificity; allowance for two missed cleavages; 20 ppm mass tolerance for MS; 0.6 Da mass tolerance for MS/MS; static modification of cysteine residues with carbamidomethylation; variable methionine oxidation; and variable modification of asparagine and glutamine deamidation. A consensus workflow, comprising the Feature Mapper and the Precursor Ion Quantifier, enabled label-free quantification and reporting of fold change in protein levels between the control and experimental sample sets, as well as calculation of p-values.

### Statistical analysis

Statistical analyses were performed by GraphPad Prism 10 (GraphPad Software, Inc.). A two-way ANOVA with a multiple comparison test was performed to assess significant differences between the groups. A significant level was set at p < 0.05 for the analysis.

## Acknowledgments

This research was funded by the Intramural Research Program of the NIH (NIEHS). We sincerely thank Dr. Perry Blackshear for his support of the research, as well as Ms. Andrea Adams and Ms. Katina Johnson for assistance with mass spectrometry sample preparation and data collection. We also value Dr. Marcos Morgan’s feedback on the manuscript.

